# USMPep: Universal Sequence Models for Major Histocompatibility Complex Binding Affinity Prediction

**DOI:** 10.1101/816546

**Authors:** Johanna Vielhaben, Markus Wenzel, Wojciech Samek, Nils Strodthoff

## Abstract

**Background:** Immunotherapy is a promising route towards personalized cancer treatment. A key algorithmic challenge in this process is to decide if a given peptide (neoepitope) binds with the major histocompatibility complex (MHC). This is an active area of research and there are many MHC binding prediction algorithms that can predict the MHC binding affinity for a given peptide to a high degree of accuracy. However, most of the state-of-the-art approaches make use of complicated training and model selection procedures, are restricted to peptides of a certain length and/or rely on heuristics.

**Results:** We put forward *USMPep*, a simple recurrent neural network that reaches state-of-the-art approaches on MHC class I binding prediction with a single, generic architecture and even a single set of hyperparameters both on IEDB benchmark datasets and on the very recent HPV dataset. Moreover, the algorithm is competitive for a single model trained from scratch, while ensembling multiple regressors and language model pretraining can still slightly improve the performance. The direct application of the approach to MHC class II binding prediction shows a solid performance despite of limited training data.

**Conclusions:** We demonstrate that competitive performance in MHC binding affinity prediction can be reached with a standard architecture and training procedure without relying on any heuristics.

## I. BACKGROUND

Immunotherapy is a promising route towards personalized cancer treatment with a variety of possible realization [see 1, for a recent review]. One path is the administration of nanoparticle vaccines customized with neoantigens. The major histocompatibility complex (MHC) plays a central role in this process as it is supposed to bind to antigens (peptides) derived from pathogens in order to display them on the surface of the cell for recognition by T-cells. There are three classes of MHC molecules, where MHC class I and II are most important due to their involvement in the targeted immune response. Due to the special nature of the MHC protein, it can bind to peptides that are potentially structurally very different from each each other. Therefore, the prediction if a certain peptide binds is a very challenging task that is, however, a crucial sub task for neoantigen identification for practical realizations of personalized immunotherapy [1].

The MHC binding prediction is a well-established problem in bioinformatics with a large number of existing algorithmic solutions. Although many of them show an excelllent performance, the present algorithms typically rely on complicated training procedures, such as pretraining on prediction tasks for related alleles or training with artificial negative peptides to achieve this performance. In addition, many of them use complicated model selection procedures to select a small number of well-performing models from potentially hundreds of trained models to eventually construct an ensemble classifier. Most of the existing approaches are restricted to peptides of fixed length, where shorter sequences are padded or longer sequences are trimmed to an appropriate length by well-motivated but still heuristic rules to identify so-called binding regions. The most prominent MHC I prediction algorithms are summarized in Tab. I. We refer to dedicated reviews for more detailed comparisons [2; 3].

**Table I:**
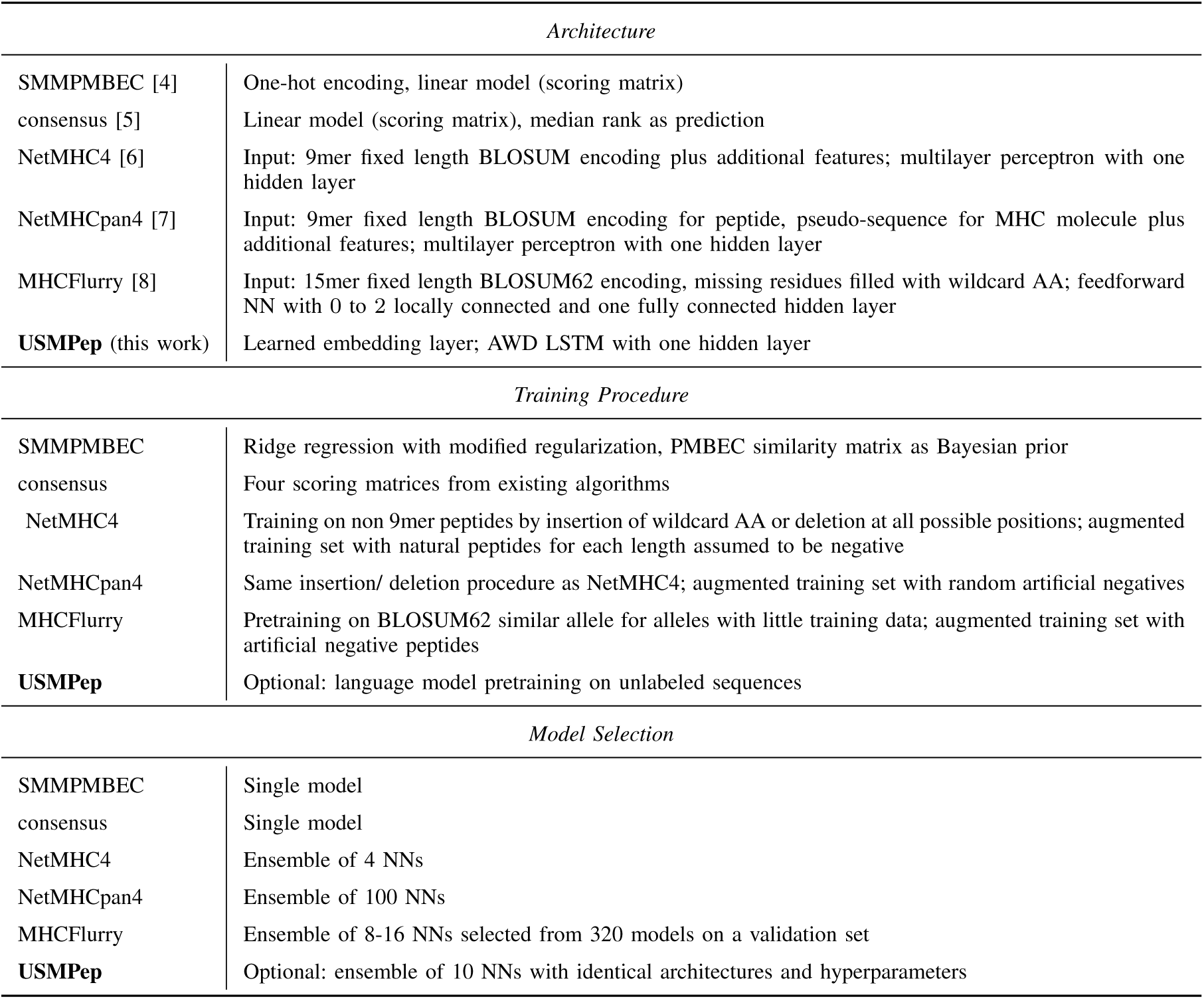
Comparison of MHC I prediction tools

Finally, not all binding prediction tools are evaluated on standard benchmark datasets, which reduces the comparability, and, even where this is the case, it is often hard to disentangle algorithmic advancements from improvements due to larger amounts of training data. In addition, statements about the generalization in the sense of the algorithm’s performance when applied to unseen data are often difficult due to potential overlaps between train and test sets, in particular as training sets often remain undisclosed. This urges for the creation for benchmark repositories, where the existing data are processed in a standardized fashion and split into training, validation and test sets.

In this manuscript, we argue that state-of-the-art performance can be reached with an astonishingly simple procedure. We use a single-layer recurrent neural network that is *trained end-to-end* on a regression task *without* any task-specific *prior knowledge* such as fixed embeddings in the form of amino acid similarity matrices. By construction, this model is able to incorporate *input of variable length without* the need for *heuristics*, such as for the identification of binding regions. The model is trained using *standard training procedures* without any artificial data or pretraining on related classification tasks. Even *single models* are is *very competitive*. Ensembling or language model pretraining only slightly improve this performance. We fix hyperparameters only once and use standard benchmark datasets to assess the model performance. We provide, amongst others, evaluation results on the recently published HPV dataset [9], demonstrating an excellent performance, which strongly suggests that the measured model performance generalizes to unseen peptide data.

Recurrent architectures have already been used previously for MHC binding prediction [10; 11] and we discuss in more detail how *USMPep* stands out from these approaches. MHCnuggets [10] is rather similar to the proposed approach (apart from the use of fixed embeddings), but relies on a complex transfer learning protocol to achieve its performance. Only limited benchmarking results are available, which makes it difficult to realistically assess its prediction performance. The very recent MHCSeqNet [11] also uses a recurrent architecture, again with pretrained rather than learned embeddings, incorporating both peptide and allele sequence to train a single prediction model for all alleles. However, the paper frames the prediction task as a classification task, which makes it difficult to align the results with the large number of existing benchmark datasets that are predominantly targeted at regression tasks. Nevertheless, the inclusion of the allele sequence represents an exciting opportunity for MHC binding affinity predicion in particular in the light of recent advances in natural language processing on tasks involving two input sequences such as question answering tasks.

## II. METHODS

### A. USMPep: Universal Sequence Models for Peptide Binding Prediction

The approach builds on the *UDSMProt*-framework [12] and related work in natural language processing [13]. We distinguish two variants of our approach, either train the regression from scratch or employ language model pretraining. A language model tries to predict the next token given the sequence up to this token, on unlabeled sequence data, here: of simulated proteasome-cleaved peptides. The architecture of the language model is at its core a recurrent neural network (LSTM) regularized by different kinds of dropout, and more specifically an AWD-LSTM model [14]. After the language model pretraining step, the model is finetuned on the regression task of MHC binding prediction by replacing the output layer with a concat pooling layer and two fully connected layers. The setup closely follows that used in [12], where protein properties were predicted. The smaller dataset sizes and shorter sequence lengths in the peptide setting (in comparison to protein classification) do not allow for building up large contexts and were accounted for by the reduction of the number of layers from 3 to 1, of the number of hidden units from 1150 to 64 and of the embedding size from 400 to 50.

Similar to [12], the training procedure included 1-cycle learning rate scheduling [15] and discriminative learning rates [13] during finetuning. Target variables for the regression model were log-transformed half-maximal inhibitory concentration (*IC*_50_)-values and a modified MSE loss function [8] that allows to incorporate qualitative data.

Dropout rate, the number of training epochs, hidden layers, hidden units and embedding dimensions, were set based on selected alleles of a particular MHC class I dataset (*Kim14* [16], see the detailed description below) by using the score on one of the provided cross-validation folds. The learning rate was determined based on range tests [15]. After this step, the aforementioned hyperparameters were kept fixed for all datasets and alleles both for MHC class I and class II prediction. In particular, neither hyperparameters nor models were selected based on test set scores.

For later convenience, the following acronyms refer to the prediction tools introduced in this work:

- **USMPep_FS_sng** single prediction model trained from scratch
- **USMPep_FS_ens** ensemble of ten prediction models trained from scratch
- **USMPep_LM_sng** single prediction model with language model pretraining
- **USMPep_LM_ens** ensemble of ten prediction models with language model pretraining

For simplicity, we consider ensembles of models with identical architectures and hyperparameters and average the final individual predictions.

### B. MHC Binding Prediction Datasets

For the downstream task of peptide MHC binding prediction, we benchmarked our model on three MHC class I and one MHC class II binding affinity datasets (details listed in Tab. II). These datasets comprise peptide sequences and the corresponding binding affinities to specific MHC alleles.

**Table II:** Details of training and test datasets. The threshold for MHC class I binders is *<* 500 nM, except for the HPV dataset, where the threshold is *<* 100 000 nM. For MHC class II binders, the threshold is *<* 1000 nM.

| Dataset | usage | total size | share of binders | # alleles | median size | share of quant. measurements | sequence length |
| --- | --- | --- | --- | --- | --- | --- | --- |
| MHC class I |  |  |  |  |  |  |  |
| *BD2009* | train | 117 326 | 0.25 | 53 | 1971 | 0.58 | 8–11 |
| *Blind* | test | 27 680 | 0.33 | 53 | 470 | 0.58 | 8–11 |
| *MHCFlurry18* | train | 120 720 | 0.25 | 32 | 3659 | 0.68 | 8–15 |
| *IEDB16_I* | test | 2827 | 0.54 | 32 | 73 | 1.0 | 9 |
| *MHCFlurry18* | train | 68 117 | 0.26 | 7 | 6884 | 0.64 | 8–15 |
| *HPV* | test | 743 | 0.34 | 7 | 125 | 0.37 | 8–11 |
| MHC class II |  |  |  |  |  |  |  |
| *Wang10* | train | 23 203 | 0.37 | 24 | 999 | 1.0 | 15–37 |
| *IEDB16_II* | test | 15 691 | 0.33 | 24 | 641 | 1.0 | 15 |

a. *Kim14:* is a commonly used binding affinity dataset compiled by [16], available on the Immune Epitope Database (IEDB)^1^ [17], and is split into a non-overlapping training (*BD2009*) and test set (*Blind*). Similar peptides (of same length with at least 80% sequence identity) shared by training and test set were removed from *Blind*. For *BD2009*, we selected the provided cross-validation split without similar peptides between the subsamples (*“cv_gs”*). There are 53 class I alleles (human and mouse/macaque alleles) with respectively 117 326 and 27 680 affinity measurements in *BD2009* and *Blind*. For comparability with recently developed systematical benchmarks [2; 9] we tested *USMPep* on two further MHC I datasets, which we refer to as *HPV* and IEDB_16. The training data of the tools reported in the literature vary in size and compilation. We trained our models on data provided by [8] and refer to this dataset as *MHCFlurry18*. It is assembled from an IEDB snapshot of December 2017 and the *Kim14* dataset.
b. *HPV:* is a recently published dataset and consists of 743 affinity measurements of peptides derived from two human paillomavirus 16 (HPV16) proteins binding to seven HLA class I alleles [9]. Peptides were considered as binders if they had *IC*_50_-values below 100 000 nM. For peptides classified as non-binders, quantitative measurements are not available.
c. *IEDB16_I:* is made up of an IEDB snapshot of October 2016 [2]. It was filtered for quantitative measurements with *IC*_50_ ≤ 50 000 nM and 9mer peptides. Training sequences of other tools were removed from the dataset. It consists of 2827 affinity measurements across 32 class I alleles. We removed any sequences occurring in the test dataset from our training data *MHCFlurry18*. In addition, we trained and tested *USMPep* on MHC class II binding data:
d. *Wang10:* is an experimental binding affinity dataset from the IEDB site^2^ based on the dataset by [18]. We used it to train our prediction tools.
e. *IEDB16_II:* is a MHC II test dataset provided by [2] from the same IEDB snapshot as the MHC I *IEDB16_I* test set above, filtered for quantitative measurements with *IC*_50_ ≤ 50 000 nM and 15mer peptides. After removing sequences present in the training data, 15 034 affinity measurements covering 24 alleles remained in the test dataset. We benchmarked our models on this dataset.

### C. Evaluation Metrics

For performance evaluation, we consider two evaluation metrics that are most frequently considered in the literature [2; 8]: The area under curve of the receiver operating statistics (AUC ROC) measures the performance of binary classifying binders and non-binders. While AUC ROC is straightforward to evaluate, it comes with the disadvantage of having to specify a threshold to turn the targets into binary labels, which discards valuable label information during the evaluation procedure. As discussed in the previous section, there exist commonly applied threshold values for the datasets under consideration but the simplicity of this procedure neglects a possible allele dependence of these threshold values [19]. This issue is circumvented by the use of ranking metrics such as Spearman *r* that evaluate the correlation between the rankings of measured and predicted affinities. Spearman *r* can only be evaluated for quantitative measurements, which discards information when evaluating on test sets containing also qualitative measurements. For both metrics, we calculated error bars based on 95% empirical bootstrap confidence intervals. For single models, we report the mean performance across 10 runs and the maximal deviation of the point estimate compared to the lower and upper bounds provided by the respective confidence intervals as a convervative error estimate.

The prediction performance across different alleles that make up a single MHC benchmark dataset can be quantified in different ways. *Overall* performance measures can be calculated across multiple alleles by concatenating all target and prediction results and evaluating the respective metrics on this set. This predominantly used but rarely discussed method has to be contrasted with reporting the *mean* or the median of the respective performance measures across all alleles, which is the default evaluation metric for related tasks such as remote homology detection [20] or transcription factor binding site prediction [21]. The difference between both evaluation approaches is related to the discussion about micro vs. macro averages for the evaluation of multi-class classification problems [22]. In particular, there are two fundamental differences between both evaluation approaches: First, the datasets enter the *overall* score with different weights determined by the size of the respective test sets, which is a weighting based on the experimental availability of binding affinities whereas the *mean* score assigns equal weight to all test sets. Second, the *overall* performance measure implicitly assumes that prediction scores are directly comparable across different alleles, which seems slightly questionable in the light of the discussion of allele-dependent binding thresholds [19]. To give the reader a complete picture of the prediction performance, we will report *overall* as well as *mean* scores. In any case, we advocate to provide individual prediction for all peptides, which allows to possibly redo the analysis using a difference performance metric at a later point in time. To this end, the peptide-wise binding affinity predictions for our tools are provided in the accompanying code repository.

## III. RESULTS

The results section is organized as follows: In Sec. III-A we present a detailed evaluation of the performance of *USMPep* for MHC class I binding affinity prediction. This is done based on three different benchmark datasets that highlight different performance characteristics. In Sec. III-B we investigate the applicability of our methods for MHC class II binding affinity prediction. Finally, we discuss language modeling on peptide data and its impact on downstream performance in Sec. III-C.

### A. MHC Class I Binding Prediction

#### 1) IEDB16 Dataset

We open the assessment of MHC class I binding prediction with results on the *IEDB16* dataset that showcases the excellent predictive performance of *USMPep*. We compare to literature results that were evaluated in a recent comprehensive benchmark [2] on this dataset. This benchmark includes evaluation metrics testing not only accuracy of binder classification, but also accuracy of binding affinity ranking and direct binding affinity prediction accuracy. Covering 32 HLA alleles, the *IEDB16* dataset reflects a broad spectrum of MHC molecules.

In Fig. 1, we show *overall* AUC ROC and *overall* Spearman *r* as reported by [2] for the latest versions of the NetMHC tools, MHCFlurry, SMMPMBEC and consensus and our scores for the different versions of *USMPep*. This is supplemented by *mean* AUC ROC and *mean* Spearman *r* compared to results provided in the data repository accompanying [2]. For the latter error bars could not be calculated for the literature approaches due to the fact that only allele-wise scores but no peptide-wise predictions were provided. In the light of the issues discussed in Sec. II-C, we advocate the use of *mean* scores rather than *overall* scores. For easy comparability, we also provide *overall* scores as they are used predominantly in the literature. It turns out that an ensemble of ten predictors with language model pretraining (*USMPep_LM_ens*), reaches the highest scores in both *mean* evaluation metrics. In this respect the results of all four *USMPep*-variants are consistent with each other and similar (within error bars) to the result of MHCFlurry, the best-performing method in the benchmark [2]. This result stresses the claims of excellent prediction performance even for a single model trained from scratch. Interestingly, the performance of all proposed prediction tools is slightly worse when considering *overall* scores. In particular, in terms of *overall* AUC ROC none of our predictors is consistent with MHCFlurry within error bars. We further investigated the origin of this performance deficiency and found that it could be traced back to a single allele, HLA-B-3801, which is peculiar in the sense that 172 of the 176 test set samples fall into a single Hobohl cluster [16] of sequences with more than 80% sequence similarity, i.e. show a particularly high sequence identity that is not seen in other test datasets. These 172 samples constitute a sizable amount of the overall 2827 test samples and strongly influence the predictive performance when using *overall* performance metrics. With this single exception in terms of *overall* ROC AUC, our proposed methods are consistent with the best-performing methods for all MHC I benchmark datasets both for *overall* and *mean* performance metrics.

**Figure 1:**
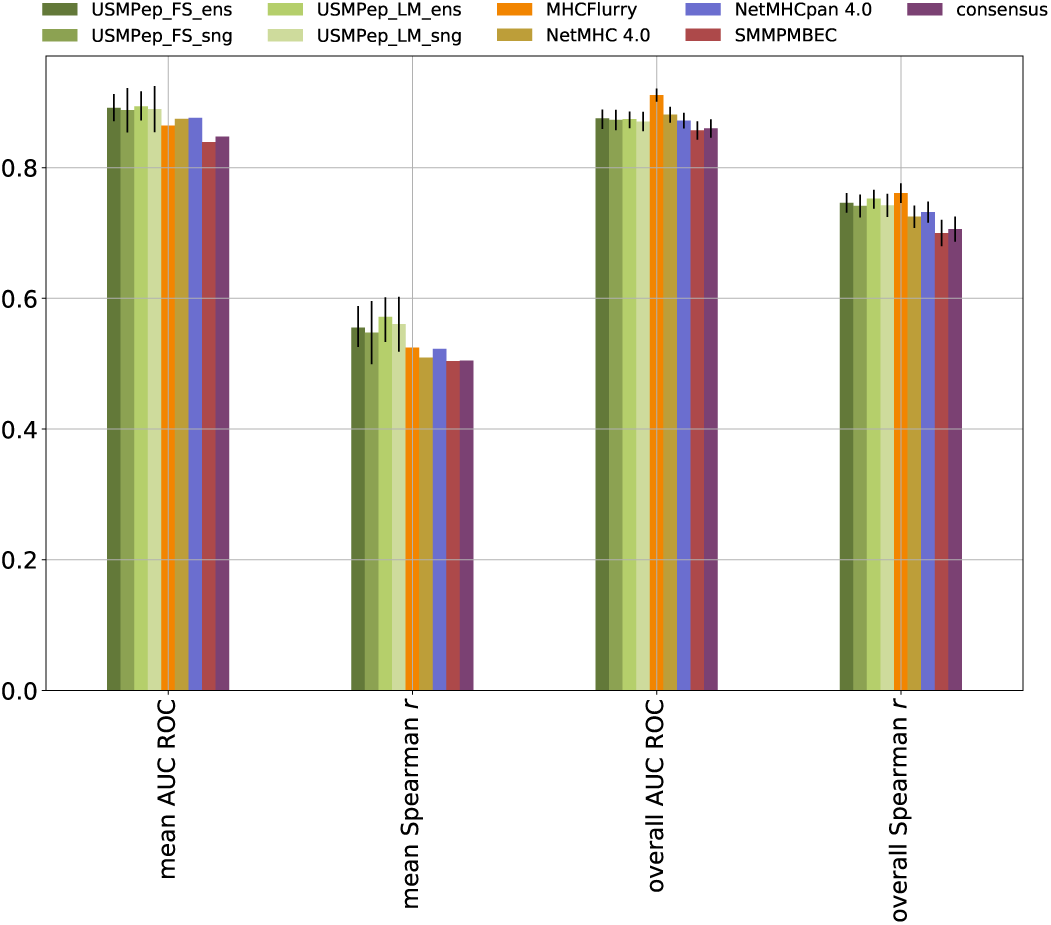
Comparison of MHC class I predictors. AUC ROC and Spearman *r* are evaluated on predictions for the *IEDB16_I* test set. AUC ROC could not be evaluated for alleles HLA-B-2704, HLA-B-1503 and HLA-B-1501, whereas Spearman *r* could not be computed for alleles HLA-B-1503 and HLA-B-1501. These alleles are therefore not included in the scores.

#### 2) HPV Dataset

As the training data is not publicly available for some MHC I prediction tools, a possible overlap between training and test datasets and correspondingly an overestimation of the predictive performance cannot be excluded. The same applies to the most common procedure of reducing the overlap between training and test set by merely removing sequences from the test set that are also contained in the training set in identical form rather than using more elaborate measures for sequence similarity. These issues can be circumvented by a performance evaluation on a dataset of different origin that has so far not been used to train MHC prediction tools. This applies to the recently released *HPV* binding affinity data [9]. However, in this benchmark, it is not possible to disentangle superior prediction performance due to larger amounts of training data from algorithmic advances since size and compilation of the training set of the algorithms vary.

As there are only quantitative measurements for the peptides considered as binders, we chose to evaluate the predictive performance only based on AUC ROC. We report the performance of all models considered in [9] and our tools measured by AUC ROC in Tab. III, where we used the predictions provided by [9]. Our *USMPep* tools show an excellent prediction performance. For three out of seven alleles an *USMPep*-model even reaches the highest AUC ROC. All neural-network-based predictors show a similar AUC ROC evaluated across all measurements in the dataset, while the ensemble with language model pretraining (*USMPep_LM_ens*) shows the highest *mean* and *overall* scores among all prediction tools. As for the *IEDB16* dataset, even the single model *USMPep*-tools are very competitive.

**Table III:**
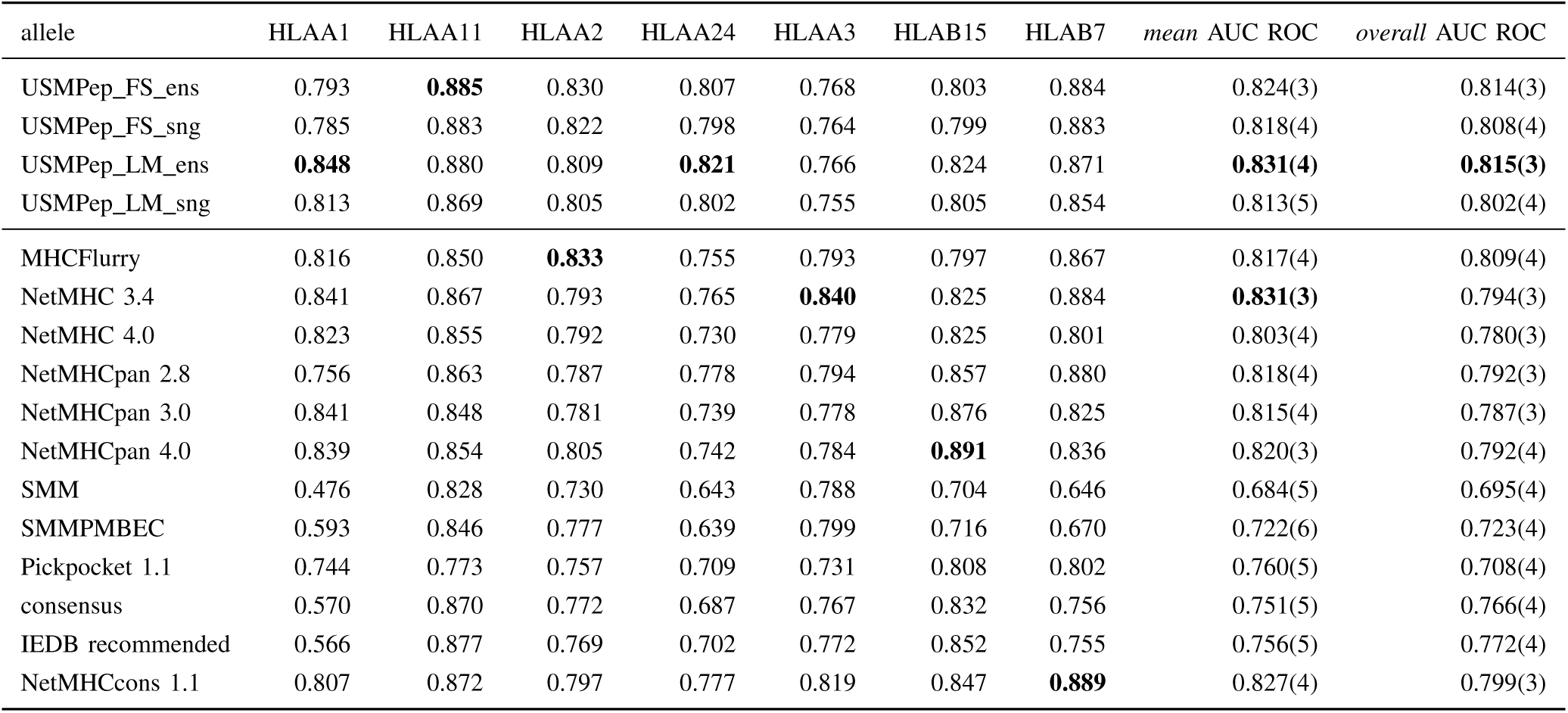
Benchmarking MHC class I predictors on recently published binding affinity data (*HPV16*). Predictive performance is evaluated by AUC ROC (threshold for binders *<* 100 000 nM) on single alleles and across all alleles (*mean* and *overall*). The scores for literature approaches were calculated based on peptide-wise predictions provided in [9].

It is instructive to investigate the performance of the different MHC prediction tools restricted to peptides of a certain length, which is only possible for the HPV dataset, where pepeptide-wise predictions for all literature approaches are provided. The result of such an analysis is shown in Fig. 2. Our tools outperform the other models on 11mer peptides. On 10mer peptides, our ensemble with language model pretraining (*USMPep_LM_ens*) and NetMHC 3.3 show a higher AUC ROC than the other tools. For 9mers our model is outperformed by most other tools and shows worse performance on 8mers. This performance gap can be explained by the fact that the internal state of the recurrent neural network has to build up over the sequence. The longer the peptide, the more context is available, which is why *USMPep* generates more accurate predictions for long sequences than for shorter ones.

**Figure 2:**
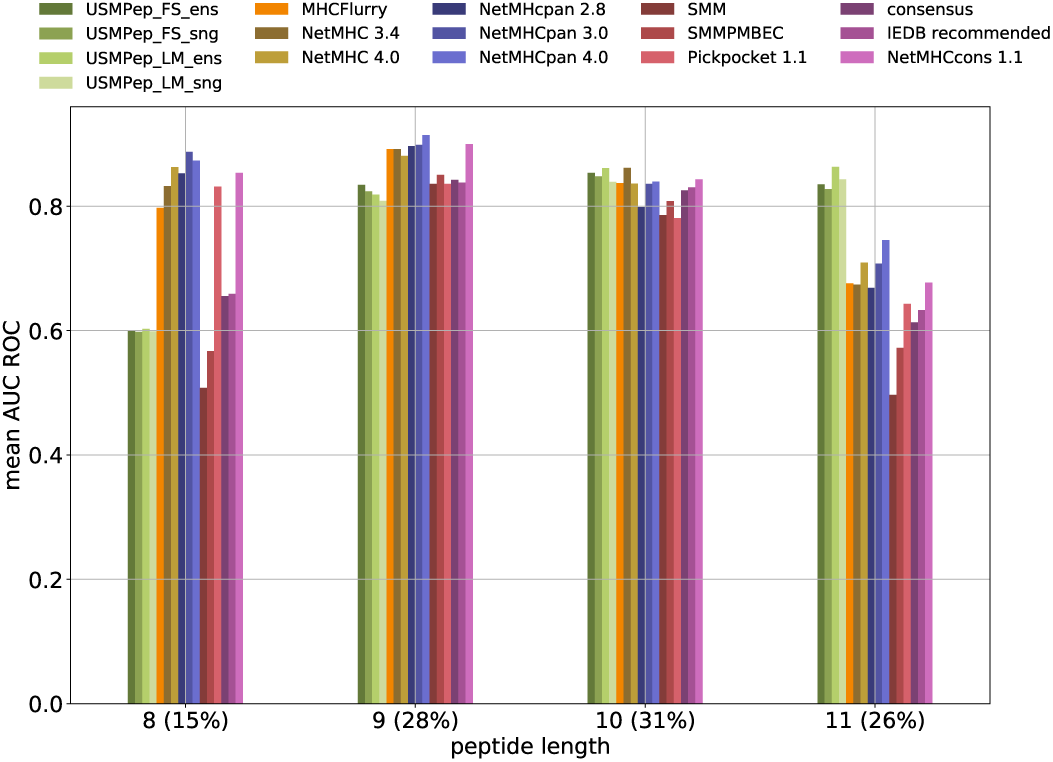
Evaluating MHC class I predictors on recently published binding affinity data (*HPV16*) grouped by peptide length. Predictive performance is evaluated by *mean* AUC ROC. For allele-wise and overall performance comparisons see Tab. III

#### 3) Kim14 Dataset

As final benchmark dataset for MHC class I prediction, we consider the *Kim14* dataset that is interesting for a number of reasons: In order to investigate how the predictive power of our approach depends on the size of the training data set, we trained and tested our model on the *Kim14 BD2009* and *Blind* data. The authors of [8] kindly provided us with the *Blind* predictions of their tool trained on *BD2009*, which allow for a direct comparison with a state-of-the-art tool. Corresponding training routines are by now also available in the code repository accompanying [8].

First, we compare the prediction success measured by AUC ROC and Spearman *r* computed across all alleles (Fig. 3). No MHCFlurry predictors exist for alleles HLA-B-2703, HLA-B-0803 and HLA-B3801 with rank 45, 49 and 52 due to insufficient training data. These alleles were therefore also excluded for the scores of our tools. The predictors perform very similarly with regard to all metrics. Our pretrained tool *USMPep_LM_ens* performs only slightly better than *USM-Pep_FS_ens* trained from scratch. This also holds for the single model versions. Both *USMPep* ensemble predictors are compatible with MHCFlurry.

**Figure 3:**
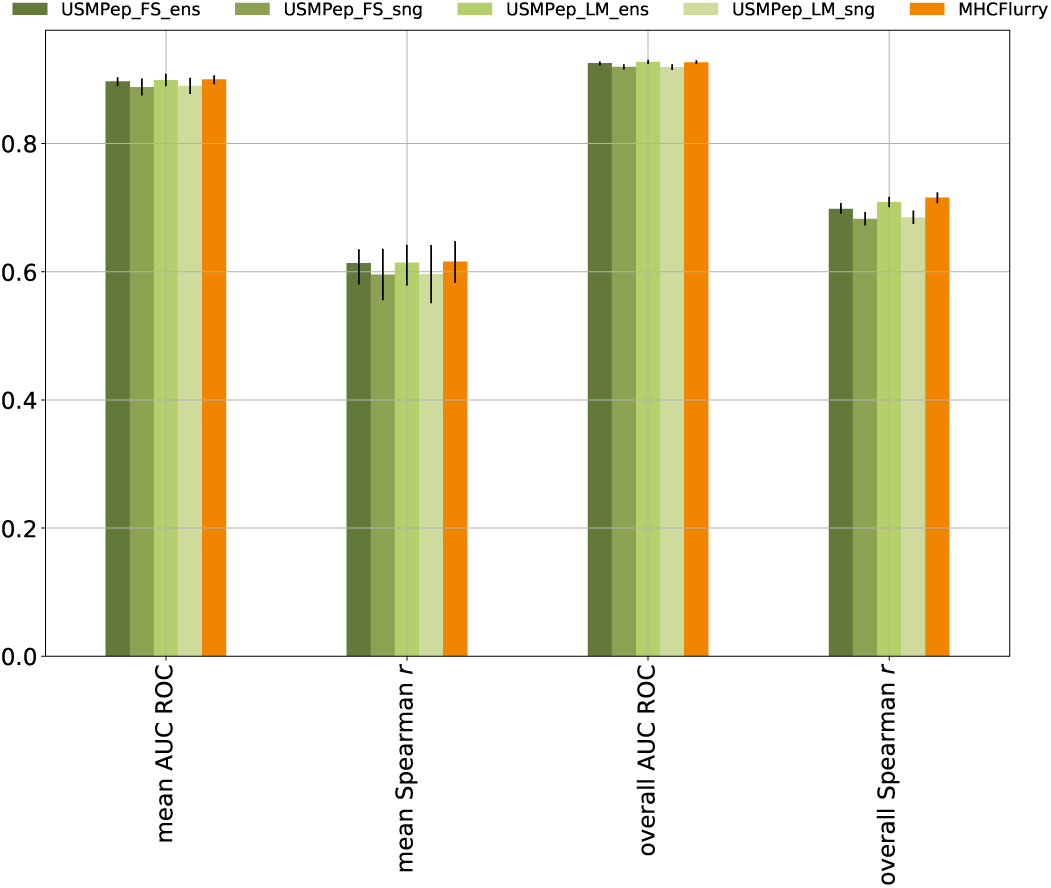
Performance of *USMPep* and MHCFlurry on MHC class I binding prediction. Both models were trained on the *Kim14 BD2009* data. AUC ROC and Spearman *r* were evaluated on the predictions for the *Blind* test set. AUC ROC could not be evaluated for allele HLA-B-4601, whereas Spearman *r* could not be computed for allele HLA-B-4601 and HLA-B-2703. These alleles are therefore not included in the scores.

Second, to examine the impact of the training set size, we report allele-wise Spearman *r* scores in Fig. 4 for our predictors and MHCFlurry. The alleles are ranked by the size of the corresponding training set. While 9528 training sequences exist for the rank 0 MHC molecule HLA-A-0201 there are only 136 training peptides for allele HLA-B-3801 with rank 52. Spearman *r* is only shown for alleles with more than 25 quantitative measurements. The allelewise performance gap between the tools becomes more pronounced the less training data are available, yet none of the models outperforms the others for the subset of alleles with less than 1000 training data points (rank 33 to 52). This is interesting considering the fact that for alleles with fewer than 1000 training measurements, MHCFlurry was pretrained on an augmented training set with measurements from BLOSUM similar alleles, *USMPep_LM_ens* was pretrained on a large corpus of unlabeled peptides and *USMPep_FS_ens* in contrast only saw the training sequences corresponding to one MHC molecule. These results stress that further efforts might be required to truly leverage the potential of unlabeled peptide data in order to observe similar improvements as seen for proteins [12] in particular for small datasets.

**Figure 4:**
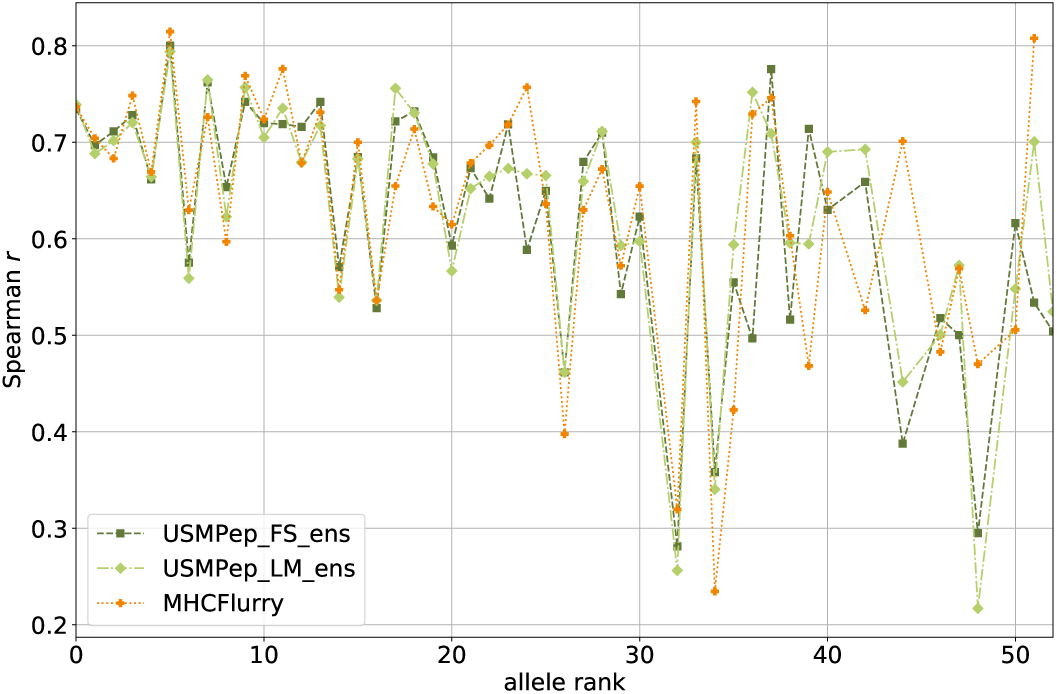
Performance of our MHC I prediction tools compared to MHCFlurry on single alleles. Spearman *r* was calculated for predictions on the *Kim14 Blind* data for alleles with more than 25 quantitative measurements. The predictors were trained on *Kim14 BD2009*. The alleles are ranked by the size of the corresponding training set (9528 peptides for rank 0 to 136 tpeptides with rank 52). No MHCFlurry predictors were provided for alleles HLA-B-2703, HLA-B-0803 and HLA-B-3801 with rank 45, 49 and 52.

### B. MHC Class II Binding Prediction

Turning to MHC Class II binding prediction, we aim to demonstrate the universality of our approach beyond its applicability to different MHC I alleles. Here, we stress again that we use the same model architecture, the same pretrained language model in case of pretraining, and even the same set of hyperparameters for all MHC class I and class II alleles. The main difference between and MHC class I and class II binding prediction is the typically larger length of 15 amino acids for MHC class II compared to at most 11 for MHC class I. The analysis of the prediction performance in dependence of the length of the peptide in the previous section suggests that this setting is particularly suitable for the *USMPep* prediction tools. Unfortunately, the reported literature results vary widely concerning the selection of training data, which makes it difficult to distinguish between algorithmic improvements and improvements due to larger amounts of training data.

The *USMPep* prediction tools, and in particular the ensemble variants, show a solid performance compared to literature results, see Fig. 5. Whereas the *USMPep*-predictors always provided the best-performing method for MHC class I prediction, it is outperformed for MHC class II by *NetMHCIIpan* and *nn_align*. We deliberately decided to train on *Wang10* instead of a more recent IEDB snapshot to work on a well-defined published dataset. However, this makes it hard to assess if the performance differences between our results and the best-performing methods can be attributed to the fact that the *USMPep*-predictors were trained using IEDB data up to 2010 whereas in particular the best-performing tools were trained on larger amounts and more recent data or if there a particular intricacies inherent to the MHC class II prediction task.

**Figure 5:**
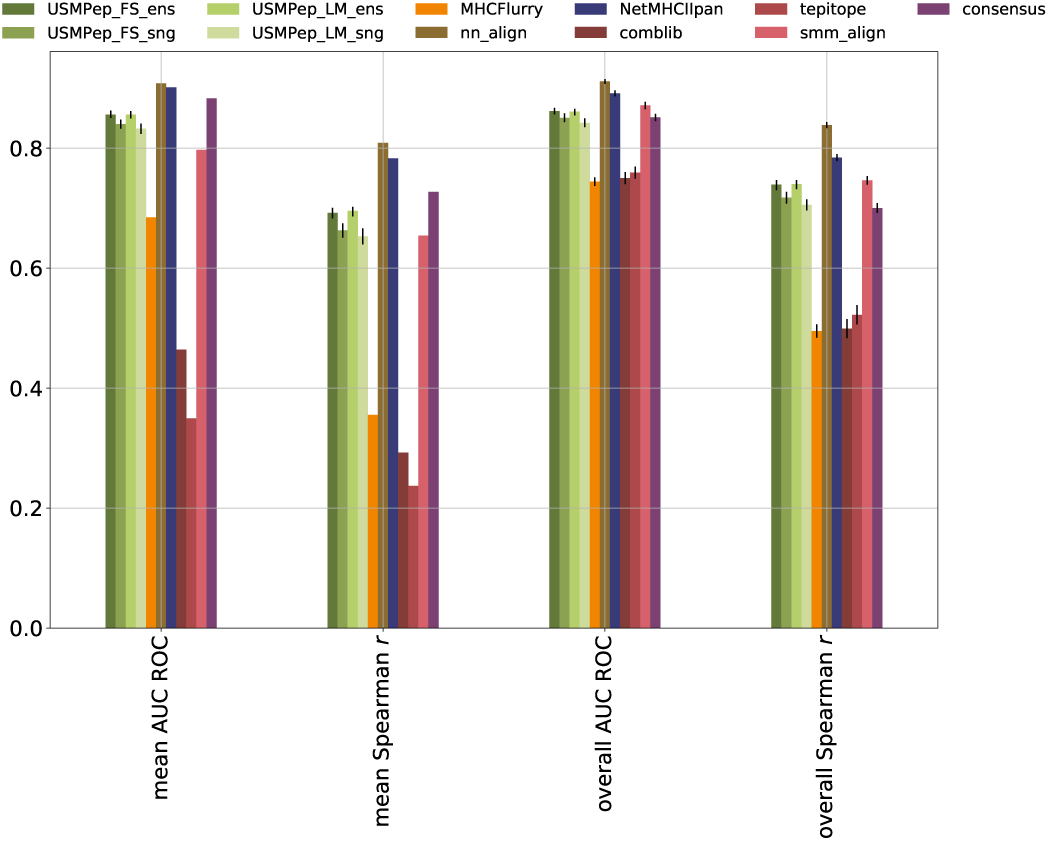
Comparison of MHC class II predictors. AUC ROC and Spearman *r* were evaluated on predictions for the *IEDB16_II* test set.

### C. Language Modeling on Peptide Data and its Impact on Downstream Performance

As final analysis, we analyze language modeling on peptide data and its impact on MHC binding affinity prediction as downstream task. To this end, we constructed a dataset of simulated proteasome-cleaved peptides to pretrain *USMpep* on a large corpus of unlabeled sequences. We filtered the SwissProt release 2018_10 for the human proteome and employed NetChop [23] to obtain proteasome cleavage sites for these proteins. The stochastic process of protein slicing was modeled by cutting with the cleavage probability provided by NetChop. We discarded sequences of less than eight and more than 20 amino acids length and obtained 6 547 641 peptides. We compare the performance of a peptide language model to that of a language model trained on human protein data using prediction accuracy as metric.

The results in terms of language model performance along with the corresponding downstream performance (MHC) on the regression task are compiled in Tab. IV and allow a number of interesting observations: First, the language model performance increases considerably when training on (proteasome-cleaved) peptide data in accordance with expectations. It is crucial to remark, that the language modeling task on peptide data poses additional difficulties compared to language modeling on protein data as the sequences are comparably short and the model thus cannot build up a lot of context. Additionally, the model does not only have to learn the normal language model task for protein data but implicitly has to learn to stochastically predict cleavage sites. Second, even we evaluated on protein data, the protein language model only reaches an accuarcy of 0.137, which is is considerably lower than the accuracy of 0.41 reported in the literature [12]. This effect is a direct consequence of the considerably smaller model size (1 instead of 3 layers; 64 instead of 1150 hidden units; embedding size of 50 instead of 400).

**Table VI:**
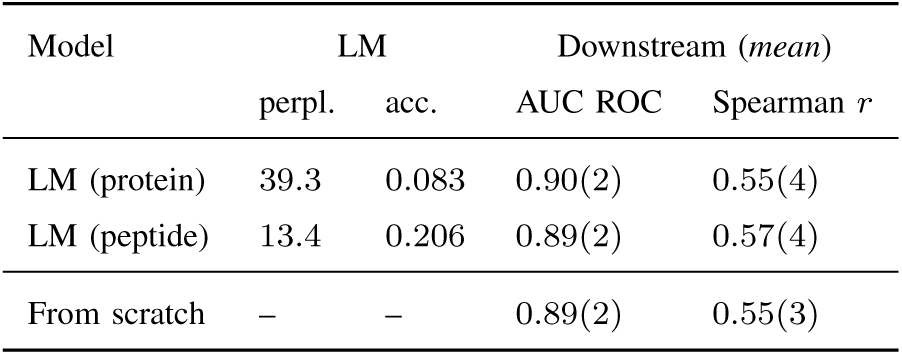
Language model and MHC class I binding affinity prediction performance. Language model metrics perplexity (perpl.) and accuracy (acc.) were in all cases evaluated on peptide data. The downstream performance corresponds to an ensemble of 10 predictors trained on the *MHCFlurry18* and evaluated on the *IEDB16_I* test set.

The details of the language model pretraining directly impact the downstream performance and show a consistent trend across all experiments described above even though the differences in downstream performance stay small and mostly remain consistent within error bars. Consistent with the general trend, the most downstream-task-adapted pretraining on peptide data performs best, generally performing slightly better than the corresponding model trained from scratch, whereas pretraining on protein data in general even leads to a loss in performance compared to training from scratch.

## IV. CONCLUSIONS

In this work, we put forward *USMPep*, a recurrent neural network that consistently shows excellent performance on three popular MHC class I binding prediction datasets as well as a solid performance on MHC class II binding prediction, see Tab. V for an executive performance summary. Most remarkably, this is achieved with a standard training procedure without incorporating artificial negative peptides, complicated transfer learning protocols or ensembling strategies and without relying on heuristics.

**Table V:** Performance summary: Rank of *USMPep* compared to competitors across the different datasets. Scores marked in bold face are best-performing or consistent with the best-performing result within error bars.

| Dataset | mean |  | overall |  |
| --- | --- | --- | --- | --- |
| | AUC ROC | Spearman $r$ | AUC ROC | Spearman $r$ |
| MHC class I |  |  |  |  |
| IEDB16_I | **1<sup>st</sup>** | **1<sup>st</sup>** | 3<sup>rd</sup> | **2<sup>nd</sup>** |
| HPV | **1<sup>st</sup>** | – | **1<sup>st</sup>** | – |
| Kim14 | **2<sup>nd</sup>** | **2<sup>nd</sup>** | **1<sup>st</sup>** | **2<sup>nd</sup>** |
| MHC class II |  |  |  |  |
| IEDB16_II | 4<sup>th</sup> | 4<sup>th</sup> | 3<sup>rd</sup> | 4<sup>th</sup> |

A central issue that prevents a true comparability of algorithmic approaches to the problem is the fact that the datasets that were used to train the prediction models differ between different literature approaches and are often not publicly available. This entangles the predictive power of a given algorithm with the data it was trained on. This urges for the creation of appropriate benchmarking repository along with standardized evaluation procedures to allow for a structured benchmarking of MHC binding prediction algorithms. As a first step, we advocate to provide binding affinity predictions for all peptides to allow fine-grained comparisons of the overall predictive performance even at a later stage as opposed to reporting just a single score summarizing the performance across all datasets.

## ACKNOWLEDGEMENTS

The source code and links to pretrained models are available at https://github.com/nstrodt/USMPep. The authors thank Patrick Wagner for discussions and work on related topics. The authors thank Timothy O’Donnell for correspondence and for kindly providing MHCFlurry predictions on the *Kim14* dataset. This work was supported by the Bundesministerium für Bildung und Forschung (BMBF) through the Berlin Big Data Center under Grant 01IS14013A and the Berlin Center for Machine Learning under Grant 01IS18037I. *USMPep* was implemented using PyTorch [24] and fast.ai [25].

## Footnotes

1 http://tools.iedb.org/main/datasets/

2 http://tools.iedb.org/mhcii/download/

## REFERENCES

[1] L. Scheetz, K. S. Park, Q. Li, P. R. Lowenstein, M. G. Castro, A. Schwendeman, and J. J. Moon, “Engineering patient-specific cancer immunotherapies,” Nature Biomedical Engineering, Aug. 2019. [Online]. Available: https://doi.org/10.1038/s41551-019-0436-x

[2] W. Zhao and X. Sher, “Systematically benchmarking peptide-MHC binding predictors: From synthetic to naturally processed epitopes,” PLOS Computational Biology, vol. 14, no. 11, p. e1006457, Nov. 2018. [Online]. Available: https://doi.org/10.1371/journal.pcbi.1006457

[3] S. Mei, F. Li, A. Leier, T. T. Marquez-Lago, K. Giam, N. P. Croft, T. Akutsu, A. I. Smith, J. Li, J. Rossjohn, A. W. Purcell, and J. Song, “A comprehensive review and performance evaluation of bioinformatics tools for HLA class I peptide-binding prediction,” Briefings in Bioinformatics, Jun. 2019. [Online]. Available: https://doi.org/10.1093/bib/bbz051

[4] Y. Kim, J. Sidney, C. Pinilla, A. Sette, and B. Peters, “Derivation of an amino acid similarity matrix for peptide:MHC binding and its application as a Bayesian prior,” BMC Bioinformatics, vol. 10, no. 1, p. 394, 2009. [Online]. Available: https://doi.org/10.1186/1471-2105-10-394

[5] M. Moutaftsi, B. Peters, V. Pasquetto, D. C. Tscharke, J. Sidney, H.-H. Bui, H. Grey, and A. Sette, “A consensus epitope prediction approach identifies the breadth of murine TCD8+-cell responses to vaccinia virus,” Nature Biotechnology, vol. 24, no. 7, pp. 817–819, 2006. [Online]. Available: https://doi.org/10.1038/nbt1215

[6] M. Andreatta and M. Nielsen, “Gapped sequence alignment using artificial neural networks: application to the MHC class I system,” Bioinformatics, vol. 32, no. 4, pp. 511–517, Oct. 2015. [Online]. Available: https://doi.org/10.1093/bioinformatics/btv639

[7] V. Jurtz, S. Paul, M. Andreatta, P. Marcatili, B. Peters, and M. Nielsen, “NetMHCpan-4.0: Improved Peptide–MHC Class I Interaction Predictions Integrating Eluted Ligand and Peptide Binding Affinity Data,” The Journal of Immunology, vol. 199, no. 9, pp. 3360–3368, 2017. [Online]. Available: https://doi.org/10.4049/jimmunol.1700893

[8] T. J. O’Donnell, A. Rubinsteyn, M. Bonsack, A. B. Riemer, U. Laserson, and J. Hammerbacher, “MHCflurry: Open-Source Class I MHC Binding Affinity Prediction,” Cell Systems, vol. 7, no. 1, pp. 129–132.e4, Jul. 2018. [Online]. Available: https://doi.org/10.1016/j.cels.2018.05.014

[9] M. Bonsack, S. Hoppe, J. Winter, D. Tichy, C. Zeller, M. D. Küpper, E. C. Schitter, R. Blatnik, and A. B. Riemer, “Performance Evaluation of MHC Class-I Binding Prediction Tools Based on an Experimentally Validated MHC–Peptide Binding Data Set,” Cancer Immunology Research, vol. 7, no. 5, pp. 719–736, Mar. 2019. [Online]. Available: https://doi.org/10.1158/2326-6066.cir-18-0584

[10] R. Bhattacharya, A. Sivakumar, C. Tokheim, V. B. Guthrie, V. Anagnostou, V. E. Velculescu, and R. Karchin, “Evaluation of machine learning methods to predict peptide binding to MHC Class I proteins,” bioRxiv, 2017. [Online]. Available: https://doi.org/10.1101/154757

[11] P. Phloyphisut, N. Pornputtapong, S. Sriswasdi, and E. Chuangsuwanich, “MHCSeqNet: a deep neural network model for universal MHC binding prediction,” BMC Bioinformatics, vol. 20, no. 1, May 2019. [Online]. Available: https://doi.org/10.1186/s12859-019-2892-4

[12] N. Strodthoff, P. Wagner, M. Wenzel, and W. Samek, “UDSMProt: Universal Deep Sequence Models for Protein Classification,” bioRxiv, 2019. [Online]. Available: https://doi.org/10.1101/704874

[13] J. Howard and S. Ruder, “Universal language model fine-tuning for text classification,” in Proceedings of the 56th Annual Meeting of the Association for Computational Linguistics (Volume 1: Long Papers). Melbourne, Australia: Association for Computational Linguistics, Jul. 2018, pp. 328–339. [Online]. Available: https://doi.org/10.18653/v1/p18-1031

[14] S. Merity, N. S. Keskar, and R. Socher, “Regularizing and optimizing LSTM language models,” arXiv preprint 1708.02182, 2017. [Online]. Available: https://www.arxiv.org/abs/1708.02182

[15] L. N. Smith, “A disciplined approach to neural network hyper-parameters: Part 1 - learning rate, batch size, momentum, and weight decay,” arXiv preprint 1803.09820, 2018. [Online]. Available: https://www.arxiv.org/abs/1803.09820

[16] Y. Kim, J. Sidney, S. Buus, A. Sette, M. Nielsen, and B. Peters, “Dataset size and composition impact the reliability of performance benchmarks for peptide-MHC binding predictions,” BMC Bioinformatics, vol. 15, no. 1, p. 241, 2014. [Online]. Available: https://doi.org/10.1186/1471-2105-15-241

[17] R. Vita, S. Mahajan, J. A. Overton, S. K. Dhanda, S. Martini, J. R. Cantrell, D. K. Wheeler, A. Sette, and B. Peters, “The Immune Epitope Database (IEDB): 2018 update,” Nucleic Acids Research, vol. 47, no. D1, pp. D339–D343, 10 2018. [Online]. Available: https://doi.org/10.1093/nar/gky1006

[18] P. Wang, J. Sidney, Y. Kim, A. Sette, O. Lund, M. Nielsen, and B. Peters, “Peptide binding predictions for HLA DR, DP and DQ molecules,” BMC Bioinformatics, vol. 11, no. 1, p. 568, 2010. [Online]. Available: https://doi.org/10.1186/1471-2105-11-568

[19] S. Paul, D. Weiskopf, M. A. Angelo, J. Sidney, B. Peters, and A. Sette, “HLA Class I Alleles Are Associated with Peptide-Binding Repertoires of Different Size, Affinity, and Immunogenicity,” The Journal of Immunology, vol. 191, no. 12, pp. 5831–5839, 2013. [Online]. Available: https://doi.org/10.4049/jimmunol.1302101

[20] J. Chen, M. Guo, X. Wang, and B. Liu, “A comprehensive review and comparison of different computational methods for protein remote homology detection,” Briefings in bioinformatics, vol. 19, no. 2, pp. 231–244, 2016. [Online]. Available: https://doi.org/10.1093/bib/bbw108

[21] B. Alipanahi, A. Delong, M. T. Weirauch, and B. J. Frey, “Predicting the sequence specificities of DNA- and RNA-binding proteins by deep learning,” Nature Biotechnology, vol. 33, no. 8, pp. 831–838, Jul. 2015. [Online]. Available: https://doi.org/10.1038/nbt.3300

[22] H. Schütze, C. D. Manning, and P. Raghavan, “Introduction to information retrieval,” in Proceedings of the international communication of association for computing machinery conference, 2008, p. 260.

[23] M. Nielsen, C. Lundegaard, O. Lund, and C. Keşmir, “The role of the proteasome in generating cytotoxic T-cell epitopes: insights obtained from improved predictions of proteasomal cleavage,” Immunogenetics, vol. 57, no. 1, pp. 33–41, Apr 2005. [Online]. Available: https://doi.org/10.1007/s00251-005-0781-7

[24] A. Paszke, S. Gross, S. Chintala, G. Chanan, E. Yang, Z. DeVito, Z. Lin, A. Desmaison, L. Antiga, and A. Lerer, “Automatic differentiation in PyTorch,” in 31st Conference on Neural Information Processing Systems (NIPS) Workshop Autodiff, 2017. [Online]. Available: https://openreview.net/pdf?id=BJJsrmfCZ

[25] J. Howard et al., “fast.ai,” https://github.com/fastai/fastai, 2018.

